# Conformational states of the cytoprotective protein Bcl-xL

**DOI:** 10.1101/2020.03.25.007740

**Authors:** P. Ryzhov, Y. Tian, Y. Yao, A. A. Bobkov, W. Im, F. M. Marassi

**Affiliations:** Sanford Burnham Prebys Medical Discovery Institute; Sanford-Burnham Medical Research Institute; Sanford Burnham Medical Research Institute; Lehigh University

## Abstract

Bcl-xL is a major inhibitor of apoptosis, a fundamental homeostatic process of programmed cell death that is highly conserved across evolution. Because it plays prominent roles in cancer, Bcl-xL is a major target for anti-cancer therapy and for studies aimed at understanding its structure and activity. While Bcl-xL is active primarily at intracellular membranes, most studies have focused on soluble forms of the protein lacking both the membrane-anchoring C-terminal tail and the intrinsically disordered loop, and this has resulted in a fragmented view of the protein’s biological activity. Here we describe how these segments affect the protein’s conformation and ligand binding activity in both its soluble and membrane-anchored states. The combined data from nuclear magnetic resonance (NMR) spectroscopy, molecular dynamics (MD) simulations, and isothermal titration calorimetry (ITC) provide information about the molecular basis for the protein’s functionality and a view of its complex molecular mechanisms.

**SIGNIFICANCE:** The human protein Bcl-xL is a key regulator of programmed cell death in health and disease. Structural studies, important for understating the molecular basis for its functions, have advanced primarily by deleting both the long disordered loop that regulates its activity and the C-terminal tail that anchors the protein to intracellular membranes Here we describe the preparation and conformations of full-length Bcl-xL in both its water-soluble and membrane-anchored states. The study provides new biophysical insights about Bcl-xL and its greater Bcl-2 protein family.

## INTRODUCTION

The functions of Bcl-xL as a major suppressor of cell death in cancer and other human diseases make it an important target for structural studies aimed at understanding its molecular mechanisms (1, 2). Bcl-xL is a member of the Bcl-2 family of apoptosis regulatory proteins (3). It localizes predominantly to the mitochondrial outer membrane (4-7), but it also appears to exist in dynamic equilibrium between cytosolic and membrane-associated populations (8, 9). Its cytoprotective activity is associated with its ability to bind cytotoxic Bcl-2 protein ligands, thereby preventing their association with mitochondria and blocking membrane permeabilization and mitochondrial destabilization (10).

The structure of Bcl-xL (11, 12) provided the framework for understanding how the Bcl-2 family proteins regulate apoptosis (1, 2). Bcl-xL shares sequence and structural similarity with other Bcl-2 family members. Eight amphipathic α-helices, spanning four Bcl-2 homology (BH) motifs, fold to form a soluble globular head domain in which a surface-exposed hydrophobic groove serves as the primary binding site for the BH3 motifs of pro-apoptotic Bcl-2 ligands. A C-terminal tail is required for both membrane association and cytoprotective activity (9). Moreover, an intrinsically disordered loop located between helices α1 and α2 carries multiple sites for posttranslational modifications – phosphorylation, deamidation and cleavage by caspases – that can dial the protein activity from cytoprotective to cytotoxic (13-34).

Structural studies have focused primarily on truncated forms of Bcl-xL lacking the C-terminal tail and/or the disordered loop. Previously, we showed (35, 36) that the tail tucks into the BH3-binding groove in the water-soluble state of the protein, while in the membrane-bound state, it forms an α-helix that anchors the protein to the lipid bilayer membrane, maintaining both the canonical structure of the head domain and its high-affinity for BH3 ligands. The roles of the disordered loop and tail in the context of wild-type, full-length Bcl-xL, however, have not been explored, and precisely how the three protein components – head, tail and loop – work together to coordinate biological function in the cytosolic and membrane-anchored states remains incompletely understood. Here we combine data from nuclear magnetic resonance (NMR) spectroscopy, molecular dynamics (MD) simulations, and isothermal titration calorimetry (ITC) to describe how the three structural elements affect the structure and activity of soluble and membrane-anchored Bcl-xL.

## MATERIALS AND METHODS

### Protein Preparation and nanodisc preparation

The BID_BH3_ peptide, corresponding to residues 80-99 in the BH3 domain of human BID, was obtained commercially (GenScript). The MSP1D1Δh5 protein for nanodisc preparation was produced in *E. coli* as described (42). Four sequences of Bcl-xL were prepared (Fig. S1). Bcl-xL-ΔLΔC (residues 1-44,85-212) was purified as described previously (35, 36). Bcl-xL-ΔL (residues 1-44,85-233), Bcl-xL-ΔC (residues 1-212), and full-length Bcl-xL (residues 1-233) were expressed and purified with an intein expression system as follows. The DNA sequences were cloned into the NdeI and SapI restriction sites of the pTYB1 vector (New England Biolabs), which results in the intein (*Saccharomyces cerevisiae* VMA1 gene) and chitin binding domain fused to the C-terminus of Bcl-xL. Plasmids were transformed in *E. coli* BL21(DE3) cells, and cells were grown in M9 minimal media containing (^15^NH_4_)_2_SO_4_ and ^13^C-glucose (Cambridge Isotope Laboratories) to obtain ^15^N/^13^C labeled protein. Cells were grown at 37°C up to OD_600_=0.8, then induced with 1 mM isopropyl 1-thio-β-D-galactopyranoside, and incubated with shaking at 18°C overnight. After harvesting (7,200 x g, 4 °C, 15 min), cells were stored overnight at −80 °C.

Cells harvested from 1000 ml of culture were suspended in 30 ml buffer A (25 mM Tris-Cl pH 8, 150 mM NaCl, 1 mM EDTA) and lysed by three passes through a French press. The soluble fraction was applied to 10 ml of chitin beads (New England Biolabs) equilibrated with chilled buffer A. After extensive washing with buffer A, 20 ml of cleavage buffer (buffer A + 30 mM dithiothreitol) were added, and the beads were incubated with gentle rocking, overnight, at 4°C. Cleaved protein was eluted from the beads with buffer A and analyzed by SDS-PAGE.

For Bcl-xL-ΔC, the eluted fraction was concentrated and further purified by size exclusion chromatography (HiLoad 16/60 Superdex 75 column; GE Healthcare) in NMR buffer (25 mM Na-phosphate pH 6.5, 1 mM EDTA, 2 mM DTT). Protein fractions were verified by SDS PAGE and then transferred into NMR buffer by centrifugal concentration (Amicon Ultra 15 concentrator with 10 kD cutoff; Millipore) and stored at 4°C.For Bcl-xL, the chitin elution fraction contained a substantial amount (∼90%) of insoluble protein. The soluble and insoluble protein fractions were separated by centrifugation (20,000 x g, 4°C, 30 min) before further purification. The soluble fraction was purified as described for Bcl-xL-ΔC. The insoluble fraction was dissolved in buffer B (25 mM Tris-Cl pH 8, 150 mM NaCl, 1 mM EDTA, 2 mM DTT, 6 M Urea, 10 mM SDS) and then purified using size exclusion chromatography (HiLoad 16/600 Superdex 200; GE Healthcare) in buffer B. Purified protein was verified by SDS PAGE, then precipitated by dialysis against water, lyophilized and stored at −20°C.

Bcl-xL and Bcl-xL-ΔL were reconstituted in nanodiscs as described (35, 36). The nanodiscs contained 2 mg of Bcl-xL and had a lipid to protein molar ratio of 1:100. The membrane was composed of a 4:1 molar mixture of dimyrystoyl-phosphatidylcholine (DMPC) and dimyrystoyl-phosphatidylglycreol (DMPG). nanodiscs were transferred to NMR buffer with an Amicon centrifugal concentrator.

### NMR experiments

Solution NMR experiments were performed on a Bruker Avance 600 MHz spectrometer with a ^1^H/^15^N/^13^C triple-resonance Bruker cryoprobe. Assignments of the N, HN, CA and CB resonances were obtained using ^1^H/^15^N/^13^C HNCA (47) and HNCACB (48) experiments. Chemical shifts were referenced to the H_2_O resonance (49). Secondary structure was characterized by analyzing the chemical shifts with TALOS+ (50, 51). Chemical shift perturbations of ^1^H and ^15^N signals (ΔHN) were calculated as: ΔHN = ½[(ΔH)^2^+(ΔN/5)^2^]. The NMR data were processed using TopSpin (Bruker) and analyzed using CCPNMR (52).

### MD simulations

All-atom MD simulations were performed using the CHARMM36 force fields for protein and lipids (53-55), with the TIP3P water model (56), as described (57). The temperature and pressure were maintained at 298.15 K and 1 bar. MD production simulations were conducted with NAMD2.11(58), using a Tesla C2075 GPU, for 400 ns (soluble state) or 350 ns (membrane state), after 20 ns of equilibration. The last 350 ns of trajectories were used for analysis, which was performed with the jupyter interface of MDanalysis 0.20.1 (59). All systems were prepared using CHARMM-GUI (60) and equilibrated with the CHARMM-GUI Membrane Builder standard protocol (61) and simulation inputs (62).

To generate the initial model of soluble Bcl-xL (Fig. S2C), the head domain was taken from the structure of the Bcl-xL/Bak-BH3 complex (12) (PDB: 1BXL), which has both tail (residues 210-233) and loop (residues 45-84) deleted. The loop (residues 21-86) was taken from the structure of tail-truncated Bcl-xL (11) (PDB: 1LXL) and the tail (residues 210-233) was inserted after R209 to replace the C-terminal His-tag. based on the structure of the isolated tail (36) (41), residues 211-230 were modeled as a helix, and docked in the BH3 groove by sequence-based alignment (Fig. S2D) with the coordinates of the Bak-BH3 helix in the complex. The model was solvated in water with 50 mM NaCl. The system for MD simulation contained 13,009 water molecules and a total of 42,645 atoms, in a 100 Å x 84 Å x 96 Å truncated octahedral box.

The initial model of membrane-anchored Bcl-xL (Fig. S2E) was derived from the structure of tail-truncated Bcl-xL (11) (PDB: 1LXL), by replacing the C-terminal His-tag with tail residues 210-233, and modeling residues 211-230 as an α-helix (36, 41). The C-terminal helix was embedded in a phospholipid bilayer composed of a 4:1 molar ratio of DMPC and DMPG, surrounded by water with 50 mM NaCl. The system for MD simulation contained 156 DMPC molecules, 39 DMPG molecules, 16,802 water molecules and a total of 76,804 atoms, in a 100 Å x 100 Å x 150 Å box.

The orientation of each helix relative to the membrane normal was obtained by calculating the geometry center (g) for the backbone heavy atoms of each overlapping set of four consecutive residues (residues i – i+3, i+1 – i+4, i+2 – i+5, …) along each helix, and then fitting them to a single vector by linear regression. For each n-residue helix there are j=n-3 overlapping four-residue segments and j corresponding g values.

### Isothermal Titration Calorimetry

Isothermal titration calorimetry experiments were performed using iTC200 instrument (Microcal) at 23 °C with 15-30 μM Bcl-xL in the ITC cell and 200-350 μM Bid_BH3_ peptide in the injection syringe, dissolved in the NMR buffer. Protein and peptide concentrations were estimated by absorbance at 280 nm. The data was processed and analyzed using ORIGIN software (Microcal). The data were fitted to a binding model corresponding to a single-site for extraction of dissociation constants (Kd). Control titrations were done by titrating against either buffer or buffer containing empty nanodiscs.

## RESULTS AND DISCUSSION

### Conformation of soluble Bcl-xL

We expressed and purified wild-type, full-length Bcl-xL using an intein-based *E. coli* expression system (37) that drives target protein production as an N-terminal fusion. Previously (36), we showed that expression systems with the reverse configuration, where Bcl-xL forms the C-terminal fusion partner, result in truncation after M218 of soluble Bcl-xL. The intein system, by contrast, yields intact, soluble protein. As noted for other expression systems, the solubility of Bcl-xL is low: *bcl-xL* gene induction produces only limited levels (∼10%) of soluble protein and Bcl-xL accumulates predominantly (90%) in the insoluble cellular fraction. As observed previously (35, 36), both soluble and insoluble fractions can be purified to obtain folded protein.

In the concentration range of ∼60-100 µM, soluble Bcl-xL is predominantly monomeric (Fig. S1) and yields well-resolved NMR spectra (Fig. 1A). This result is in line with the very high (∼600 µM) dissociation constant for dimer formation that has been reported based on fluorescence quenching experiments (38). In cells, soluble Bcl-xL has been proposed to form dimers in which the tail of one monomer associates with the BH3-binding grove of the other (38, 39). In vitro, however, the ^1^H/^15^N NMR spectra show no evidence of such dimerization. Limited protein solubility and low NMR intensity precluded complete resonance assignment and structure determination. The absence of tail signals reflects the presence of amide hydrogen exchange with water and/or conformational dynamics in the μsec-msec time scale for this region of the protein. At 25°C, the spectra are dominated by signals from the α1-α2 loop, while raising the temperature to 45°C caused most tail peaks to disappear.

**Figure 1.**
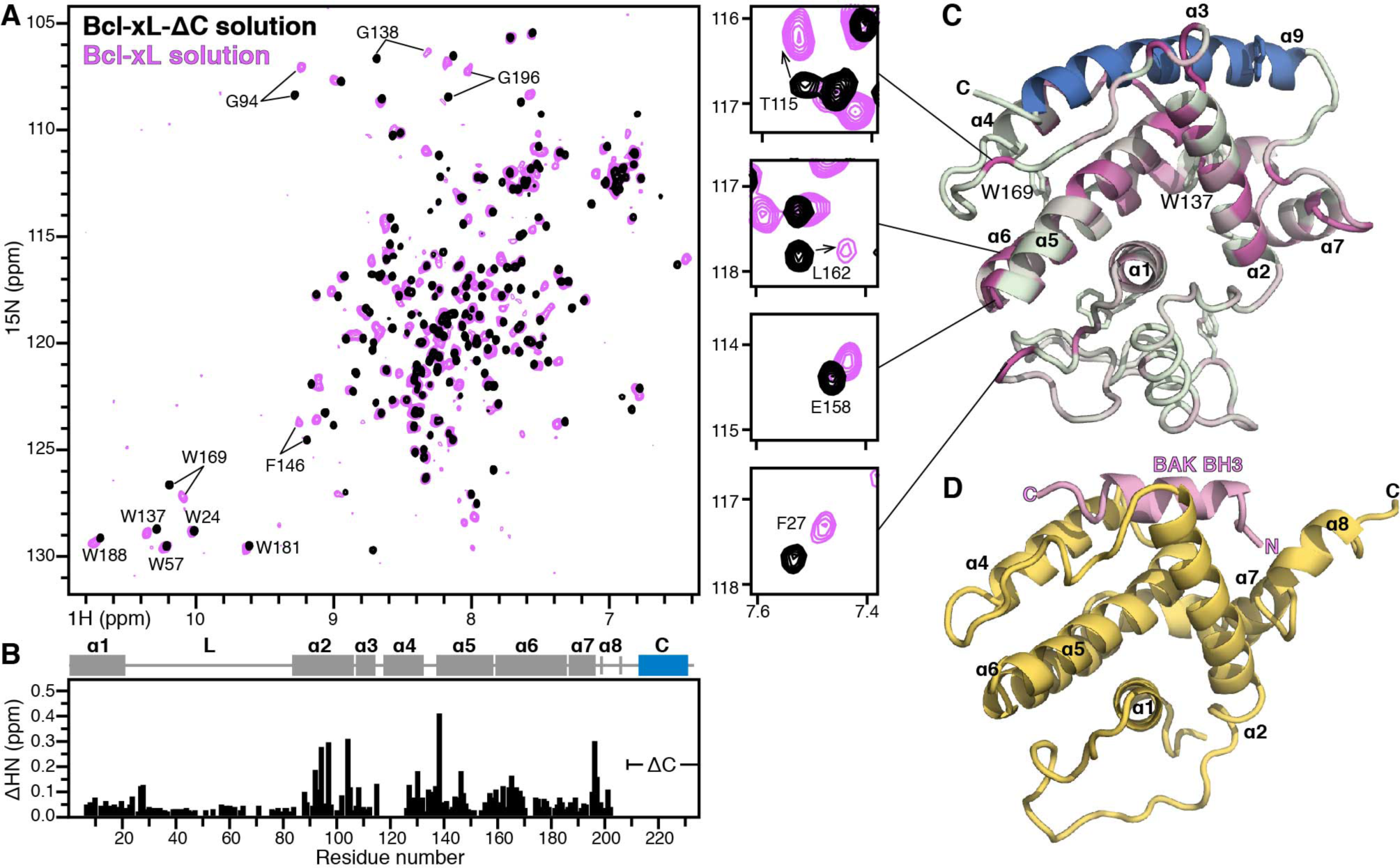
Conformation of soluble full-length Bcl-xL. **(A)** ^1^H/^15^N HSQC NMR spectra of ^15^N labeleld Bcl-xL ΔCt (black) and wild-type Bcl-xL (mauve). The spectra were recorded at 45°C. Selected regions are expanded to highlight specific perturbation sites. **(B)** Profile of tail-induced chemical shift perturbations across the sequence of Bcl-xL. Bars represent the combined difference (ΔHN) of amide ^1^H and ^15^N chemical shifts, (ΔHN = [(ΔH)^2^ + (ΔN/5)^2^]^1/2^). Helix boundaries are taken from the original structures of Bcl-xL (11, 12). **(C)** Structural model of soluble Bcl-xL taken after 400-ns MD simulation. Colors reflect the magnitude of ΔHN from 0 ppm (gray) to the maximum value (0.39 ppm, mauve). The tail is colored blue. **(D)** Structure (PDB; 1BXL) of the tail- and loop-truncated Bcl-xL (gray) bound to a Bak-BH3 peptide (pink) (12).

Overall, the ^1^H/^15^N NMR spectrum of full-length Bcl-xL is similar to that of tail-truncated protein (Bcl-xL-ΔC), indicating that the tail does not disrupt the general three-dimensional fold. Notably however, distinct differences are also observed. The chemical shift perturbation profile (Fig. 1B) maps predominantly to the BH3-binding groove, covering its entire topological length, from α7-α8 at the top of the groove to the start of α6 at the bottom. The perturbation profile is distinct from that of the well-known, high-affinity association of the groove with BH3 peptides, which bind with their N-terminus near the start of α4, as illustrated by the complex of loop- and tail-truncated Bcl-xL with a Bak BH3 peptide (12) (Fig. 1D).

Perturbations of the NH signals from G94, G138 and F146 are sentinels of the bound BH3 groove, while the G196 perturbation is specific for the tail-groove interaction (35, 36). Perturbations at W169 and sites near the start of α6 reflect a more extensive engagement of the groove and are a hallmark of Bcl-xL association with longer BH3 peptides (40), but not observed for M218-truncated Bcl-xL. Finally, while the effects of the tail localize primarily to the hydrophobic BH3-binding groove, some perturbations are also observed in the α1-α2 loop, at the polar opposite end of the protein. We conclude that the conformational rearrangements associated with the tail-groove interaction are relayed to the loop, and therefore, that the loop is not totally conformationally disconnected from the head.

The NMR data reflect a protein conformation in which the tail is predominantly associated with the BH3-binding grove. To gain molecular insights about this cytosolic state we generated a model of the protein based on the NMR data, and the existing structural information, and performed all-atom MD simulations in aqueous solution. The starting model (Fig. S2C) was templated from the structure of the Bcl-xL/Bak-BH3 complex (12), which has both tail (residues 210-233) and loop (residues 45-84) deleted, while the random coil conformation for the loop was taken from the structure of tail-truncated Bcl-xL (11). Guided by the experimental NMR structure of the isolated tail peptide (36, 41), we modeled residues 211-230 as a helix and placed it in the BH3 groove by sequence-based alignment with the Bak-BH3 helix in the complex.

After 400-ns MD simulation (Fig. 1C), the tail forms a helix spanning residues 207-230 and remains associates with the BH3 pocket. As observed for the Bak-BH3 peptide complex (Fig. 1D), the segment corresponding to α3 is unraveled as the BH3 groove adjusts to accommodate its ligand. Unlike the complex, however, α8 (residues 198-205) also unravels to enable the tail to adjust its position in the groove. The experimental chemical shift perturbations for this region of the protein are substantial, and parallel the conformational change produced by the MD simulation. Notably, the loop undergoes a dramatic condensation from its initial, extended structure, and packs against α1 in the final conformation (Fig. 1C, S2). The MD simulation also produces a propensity for helical structure in the second half of the loop (residues 66-78). Helical tendency in this region is also observed in the NMR structure of Bcl-xL-ΔC (11). This region coincides with the peptide predicted to be excised upon cleavage with caspase 3 (16, 17), raising the intriguing possibility that it plays a functional role in apoptosis regulation.

Within the limitations that our starting model for MD is not an experimentally determined structure of full-length Bcl-xL, and that the MD trajectory spans a limited time scale of 400 ns, the resulting conformation is consistent with the NMR data and provides molecular insights about the cytosolic state of Bcl-xL. When viewed in the frame of the MD structural model, the ^1^H/^15^N NMR data show how the tail-groove interaction propagates allosterically from one end of the molecule to helix α1 and the loop, at the polar opposite end, such that association of the tail C-terminus with the α3-α4 turn (T115) is sensed by α5-α6 sites (L162, E158, W169) one level below, and relayed to loop sites (F27) two levels below.

### Membrane-anchored Bcl-xL

To examine the effects of the loop and tail on its membrane-inserted state, we reconstituted full-length Bcl-xL in lipid bilayer nanodiscs, prepared with a 4/1 molar mixture of the phospholipids dimyristoyl-phosphatidyl-choline (DMPC) and dimyristoyl-phosphatidyl-glycerol (DMPG), and the short membrane scaffold protein MSP1D1Δh5 (42). The NMR spectra of full-length and loop-deleted (Bcl-xL-ΔL) protein both reflect the membrane-anchored conformation (35, 36), but many chemical shift differences are also apparent (Fig. S3). As expected, marked differences map to the loop excision sites of Bcl-xL-ΔL (residues E44 and A85), but many prominent changes are also observed at more distal sites, specifically, residues 23-28 at the start of the loop, as well as α1, α2, α3, α7 and α8. The effect of the loop on the head is similar to that reported for soluble tail-truncated protein (34) where the loop was shown to cause a subtle repositioning of α3, indicating that similar head-loop contacts are present in the membrane-anchored state of the protein.

Comparison of the NMR spectra from soluble Bcl-xL-ΔC and membrane-anchored Bcl-xL reveals the effects of the membrane on the protein (Fig. 2). The chemical shift differences reflect protein-membrane interactions as well as conformational rearrangements in the linker connecting the helical transmembrane tail to α7-α8. Membrane anchoring induces prominent perturbations near the BH3-binding groove. To gain molecular insights we performed all-atom MD simulations of the protein in a lipid bilayer membrane with a similar 4/1 molar composition of DMPC/DMPG as the experimental nanodiscs. The starting structure was generated by modeling tail residues 211-230 as a helix (36, 41), then appending them to the structure of tail-truncated Bcl-xL (11) and positioning the tail straight across the lipid bilayer (Fig. S2E). After 350-ns MD simulation, the protein adopts a preferred orientation in the membrane (Fig. 2C, 3A, 3B). The tail helix spans residues 207-230 and adopts a marked tilt of ∼36° relative to the axis normal to the membrane plane. The head also adopts a preferred average orientation that places the BH3-binding groove near the membrane surface (Fig. 3C) while keeping it accessible to the aqueous milieu, consistent with the experimental chemical shift perturbations observed for the membrane-anchored protein and with the high affinity for BH3 ligands of the membrane-anchored state (35, 36, 41).

**Figure 2.**
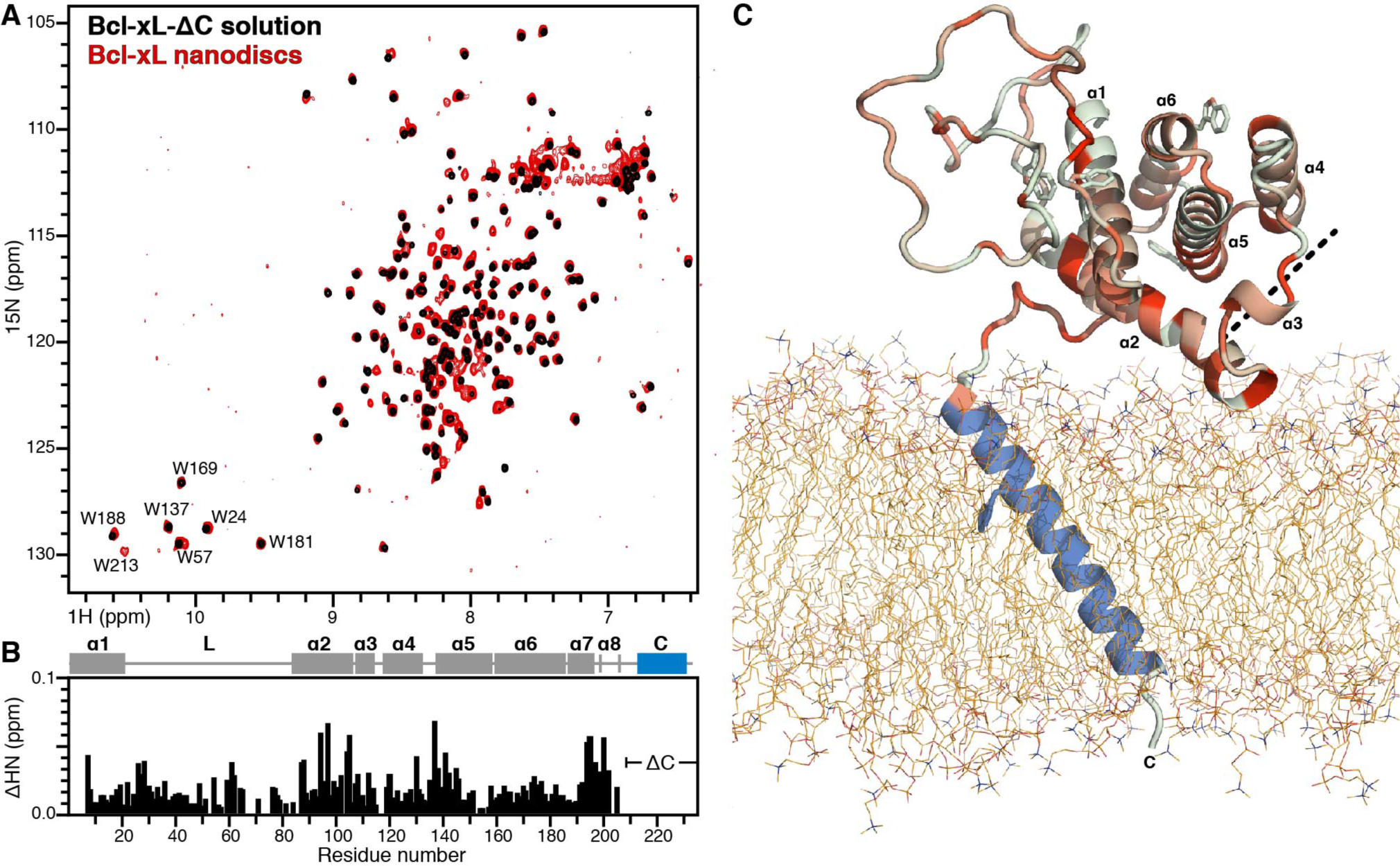
Effect of membrane-anchoring on the head and loop of Bcl-xL. **(A)** ^1^H/^15^N HSQC NMR spectra of ^15^N labeleld Bcl-xL ΔCt (black) and wild-type Bcl-xL (mauve) recorded at 45°C. **(B)** Profile of tail-induced chemical shift perturbations across the sequence of Bcl-xL. Bars represent the combined difference (ΔHN) of amide ^1^H and ^15^N chemical shifts, (ΔHN = [(ΔH)^2^ + (ΔN/5)^2^]^1/2^). Helix boundaries are taken from the original structures of Bcl-xL (11, 12). **(C)** Structural model of soluble Bcl-xL taken after 350-ns MD simulation. Colors reflect the magnitude of Δδ from 0 ppm (gray) to the maximum value (0.07 ppm, mauve). The tail is colored blue. The dashed line marks the position of the BH3-binding groove.

**Figure 3.**
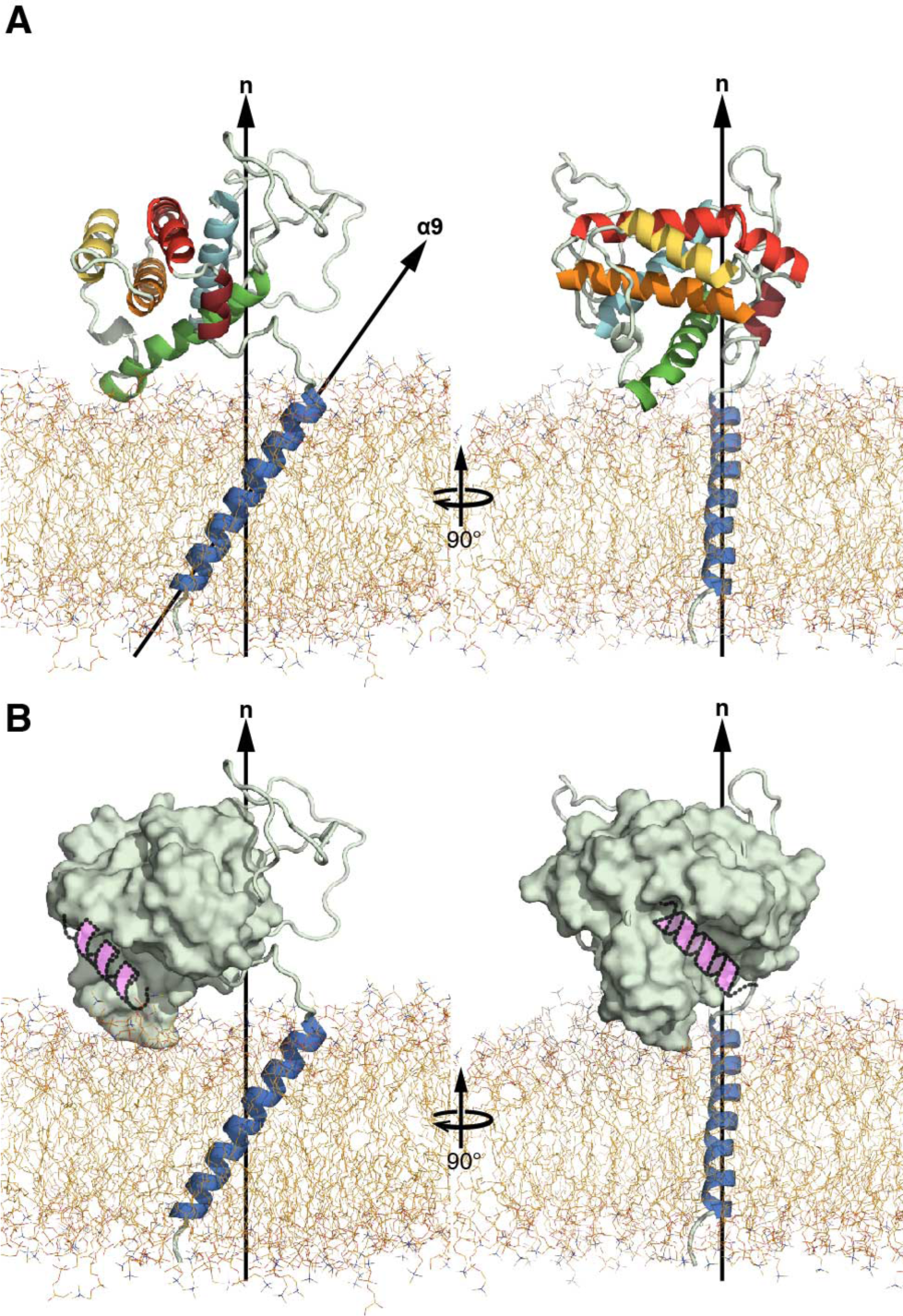
Position of membrane-anchored Bcl-xL in the membrane obtained after 350 ns MD simulation. **(A)** Orthogonal views of the protein colored by helix: α1 (cyan), α2 (green), α4 (yellow), α5 (orange), α6 (red), α7 (rust, and α9 (blue). Arrows denote the axis normal to the membrane plane (n) or the C-terminal helix axis (α9). **(B)** Orthogonal views showing the head (gray surface), tail (blue) and loop (cartoon) in the frame of the membrane. A BH3 peptide model helix (pink) is drawn into the groove to highlight its position on the head. The head adopts a preferred orientation with the BH3-binding groove near the membrane.

### Protein dynamics

NMR peak intensities provide a measure of a protein’s propensity for chemical exchange and/or conformational flexibility. The soluble and membrane-anchored states of Bcl-xL exhibit generally similar ^1^H/^15^N peak intensity profiles (Fig. 4A). In both cases, the highest intensity sites coincide with the loop, in line with the absence of both electron density in the crystal structure and medium-to long-range NOEs in the NMR spectra for this region of the protein (11). In the case of soluble Bcl-xL, higher intensity is also apparent for the α3-α4 and α7-α8 regions, in line with the lack of structure for residues 108-111 (α3), and helix unfolding for residues 199-205 (α8) to accommodate the association of the tail into the BH3 groove. For membrane-anchored Bcl-xL, higher intensity for residues 199-205 reflects the formation of a flexible linker that enables the head to reorient freely relative to the nanodisc membrane and allows the solution NMR spectrum to be detected. In both states, the intensity profile peaks at E39, S62 and R78, and exhibits distinct reduction around A50 and V65. The first half of the loop contains multiple negatively charged Glu and Asp residues whose mutually repulsive interactions may contribute to disorder and flexibility.

**Figure 4.**
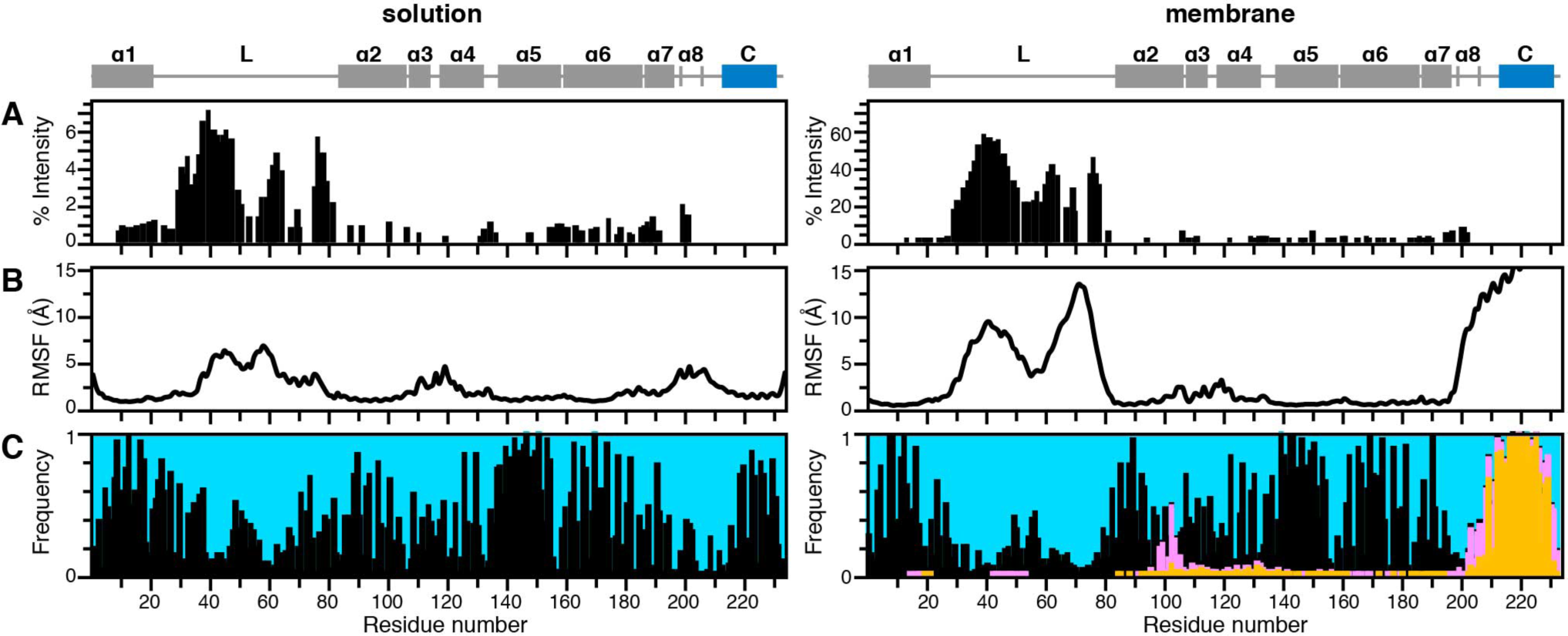
NMR Peak intensity, MD flexibility and MD contact profiles of soluble (left) and membrane-anchored (right) Bcl-xL. **(A)** Percent ^1^H/^15^N HSQC peak intensities, normalized to the signal from G21 at 1%. **(B)** Time-averaged RMSF calculated for heavy atoms over the last 300 ns of MD trajectories. Protein alignment was relative to the head (residues 1-22 and 82-196). **(C)** Interaction profile of protein residues with their environment. The bars represent frequency of occurrence within 4 Å of water (blue), phospholipid head groups (pink) or tails (gold), or protein sites (black).

The intensity profile is more extremely defined for the membrane-anchored state: when each plot is internally normalized relative to the signal intensity from G24, a well resolved signal that can be measured with high accuracy, the maximum intensity of the membrane state is 10 times greater than that of the soluble state. We attribute this effect to the substantially slower overall tumbling rate of the nanodisc assembly (35), which causes general broadening and intensity reduction of NMR signals from the tail and head domains.

It is, nevertheless, also possible that increased loop flexibility and/or hydrogen exchange rates are present in membrane-anchored Bcl-xL and contribute to this effect. The MD simulations provide some insight in this regard. The time-averaged root mean-square fluctuations (RMSF) calculated for heavy atoms over the last 300 ns of MD trajectories (Fig. 4B) parallel the experimental intensity profiles. As observed experimentally, both states have reduced fluctuations in the middle of the loop but the loop in the membrane-anchored state is twice as flexible as the soluble state. Consistent with this observation, the conformation of the loop is highly compacted around α1 in solution and somewhat more expanded in the membrane. For membrane-anchored Bcl-xL, the very high RMSF of residues 199-233 parallels the unraveling of α8, and the motional decoupling of head and tail dynamics on the nanosecond time scale that is observed experimentally by NMR (35)

The profile of intramolecular protein-protein, protein-water and protein-lipid contact frequencies (Fig. 4C) further reveals that residues in the middle of the loop make frequent contact with protein sites in the folded head domain, providing an explanation for their reduced flexibility compared to the rest of the loop. This is in line with the observation of ^1^H-^1^H NOE cross-peaks between the folded head and the W57 side chain in a deamidation mimic loop mutant of Bcl-xL (34). In the membrane-anchored state this region of the loop also has some, albeit infrequent, encounters with the membrane surface, suggesting that fluctuations between contacts with the membrane and contacts with other protein sites could contribute to enhanced loop dynamics. The contact map of membrane-anchored-Bcl-xL also reflects substantial contacts between the membrane surface and sites in the BH3-binding groove, particularly α2-α4, in line with the experimental membrane-induced chemical shift perturbations (Fig. 2) and the preferred orientation of the head domain (Fig. 3).

### The loop and tail modulate the ligand binding activity of Bcl-xL

To examine the effects of the loop and tail on the BH3 binding characteristics of Bcl-xL, we performed ITC experiments with a peptide (Bid_BH3_) spanning residues 80-99 of the Bid BH3 motif. All cases reflect the 1/1 binding stoichiometry documented for the interactions of BH3 ligands with cytoprotective Bcl-2 proteins. The binding affinity (Fig. 5A) of soluble full-length Bcl-xL for Bid_BH3_ is reduced by a factor of ∼25 relative to either the isolated head domain (Bcl-xL-ΔLΔC) or the combined loop-head domains (Bcl-xL-ΔC). This is in line with the structural model for cytosolic Bcl-xL in which the tail associates with the BH3-binding groove, restricting access by other extramolecular ligands. The data are consistent with a BH3 binding event that involves competition with the protein’s C-terminus.

**Figure 5.**
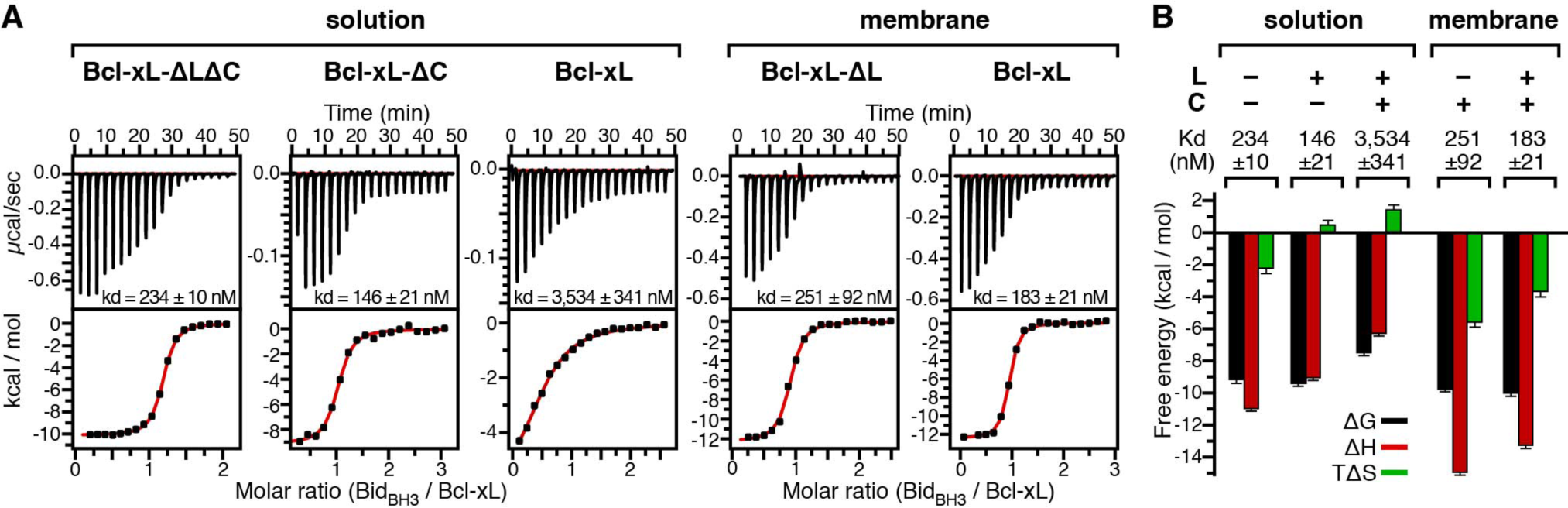
ITC titrations of BID_BH3_ peptide into Bcl-xL. **(A)** Representative calorimetric data (top) and integrated heat (bottom) are shown as functions of BID_BH3_/Bcl-xL molar ratio. The data were corrected for non-specific binding by subtracting control ITC titrations performed by titrating peptide into buffer or empty nanodiscs without Bcl-xL. Solid lines (red) are the best fits of the binding isotherms to a single-site binding model, used to extract the values of the dissociation constant (K_d_). **(B)** Plot of the thermodynamic parameters for each calorimetric titration. Values represent the average of triplicate or duplicate titrations and erroro bars represent standard deviation.

In all three cases – Bcl-xL-ΔLΔC, Bcl-xL-ΔC, and Bcl-xL – the thermodynamic binding parameters (Fig. 5B) are dominated by favorable enthalpy (ΔH), consistent with the formation of residue-specific interactions between the groove and the BH3 ligand. While the BH3 binding interaction with Bcl-xL-ΔLΔC is purely enthalpy-driven and carries an unfavorable entropic factor associated with Bid_BH3_ helix formation from an unfolded state (40), BH3 binding to full-length Bcl-xL has a substantial favorable entropic component. Displacement of the tail by the BH3 ligand likely contributes more than enough entropy gain to compensate for the entropy lost upon BH3 peptide folding.

Interestingly, reintroduction of the loop in Bcl-xL-ΔC contributes favorably to BH3 ligand affinity, enhancing it by a factor of ∼1.6 relative to Bcl-xL-ΔLΔC. BH3 ligand binding in this case is accompanied by a favorable entropy increase (∼2 kcal/mol). A parallel effect is observed for the BH3-binding signature of membrane-anchored Bcl-xL (Fig. 5B). Here, the tail is membrane-inserted leaving unrestricted access to the groove. In addition, the loop contributes a positive entropic factor (∼1.8 kcal/mol) to the binding free energy, enhancing the affinity by a factor of ∼1.4 compared to Bcl-xL-ΔL.

These results reflect a coupling of the loop with the head in both the soluble and membrane-anchored states of Bcl-xL. The data indicate that the tail-groove interaction causes subtle conformational rearrangements that transmit allosterically to the distal loop, and that the loop must interact appreciably with the head to sense these perturbations. These findings complement those of another recent study (34) where transient loop-head interactions were shown to induce a subtle repositioning of sites at the rim of the distal BH3-binding groove.

## CONCLUSIONS

Structural studies of Bcl-xL have focused primarily of tail-truncated constructs, due to the technical difficulties associated with full-length protein preparation and purification. Here we described the production of sufficient quantities of intact, wild-type Bcl-xL for NMR structural studies of either the soluble or membrane-anchored protein states. The results show that association of the tail with the BH3-binding groove reduces the affinity of Bcl-xL for a Bid BH3 ligand by a factor of ∼25. Affinity reduction due to occlusion of the groove by the C-terminal tail has been reported for Bcl-w (43, 44) and for forms of Bcl-xL encompassing part of the tail (35, 36, 45). Compared to these, the ∼25-fold effect observed for the complete tail of Bcl-xL is the most dramatic, indicating that the full inhibitory effect requires the complete length of C-terminal tail, in the case of Bcl-xL. As noted previously (35), and contrary to studies where detergent micelles were used as membrane mimics, the Bcl-xL head domain adopts the same overall structure in either its soluble or membrane-anchored state, and retains similar BH3-binding properties. The present data further illustrate the subtle contributions of the loop to the structure, dynamics and activity of Bcl-xL.

Bcl-xL like its Bcl-2 relatives that possess a membrane-anchoring tail localize predominantly to intracellular membranes. Endogenous Bcl-xL is integral to the mitochondrial outer membrane (46) and the C-terminal tail is essential for membrane integration. The conformation of membrane-anchored Bcl-xL provides a view of this predominant state of the proteins. Bcl-xL has also been reported to translocate between cytosolic and membrane-anchored states. How might this occur? The water solubility of wild-type, full-length Bcl-xL is highly curtailed, raising the question whether additional factors assist its shuttling between cellular compartments. Previously, we showed that the caspase-8 cleavage product of Bid is capable of associating with phospholipids to form nanometer size, lipoprotein particles, that are soluble and retain binding affinity for the anti-apoptotic protein Bcl-xL. This points to a potential role of lipids in mediating Bcl-2 protein mobility and interactions, and the notion that lipid-assisted cytoplasmic solubility may be important for Bcl-2 protein function. Rather than adopting exclusively lipid-free or membrane-anchored states, Bcl-xL and its Bcl-2 relatives may lead less binary lifestyles, with lipids as key partners. Whether this is true for Bcl-xL will have to be examined with additional studies aimed at resolving the protein structures and interactions in samples that closely resemble the native environment.

## SUPPORTING MATERIAL

Supporting Material can be found online.

## AUTHOR CONTRIBUTIONS

PR, YT, YY, AAB performed experiments.

PR, YT, YY, WI, and FMM analyzed data.

FMM and PR wrote the manuscript.

FMM designed the study

## ACKNOWLEDGEMENTS

This study was supported by grants from the National Institutes of Health (CA 179087 and GM 118186 to FMM) and the National Science Foundation (MCB-1727508, MCB-1810695 and XSEDE MCB-070009 to WI). It utilized the Structural Biology Resource supported by grant P30 CA030199.

